# Unlocking karst biodiversity with eDNA: a validated qPCR toolkit for discovery and monitoring the world’s largest cavefish radiation (Cyprinidae: *Sinocyclocheilus*)

**DOI:** 10.64898/2025.12.10.692593

**Authors:** Yanming Peng, Shenjie Yu, Rongjiao Chen, Yewei Liu, Tingru Mao, Ting Lin, Dan Sun, Marcio R. Pie, Jian Yang, Madhava Meegaskumbura

**Author notes:** Author for Correspondence: MM; E-mail –.

## Abstract

Environmental DNA (eDNA) enables non-invasive detection of aquatic species through trace genetic material, but uncertainties regarding DNA persistence, detection accuracy, and the limited taxonomic resolution of stygian fauna have constrained its use in subterranean aquatic environments. The species-rich cavefish genus *Sinocyclocheilus*, endemic to the karst landscapes of southwestern China, offers an excellent model for understanding these issues. With over eighty species, many of which are endemics, and many still being described, this lineage is taxonomically rich, poorly understood, and highly threatened. However, access to these deep habitats remains challenging, making eDNA detection one of the most practical methods for documenting and monitoring them. We developed and validated a qPCR assay targeting a hypervariable region of the mitochondrial 16S rRNA gene, allowing for genus-specific detection and species-level identification. Laboratory and field evaluations involving 47 representative species confirmed high specificity and a detection limit of around 20 DNA copies per reaction (R² = 0.995). Controlled degradation studies indicated that temperature and pH strongly affect DNA persistence. Field surveys across 47 karst sites (37 caves and 10 surface sites) detected *Sinocyclocheilus* eDNA in only 33 caves. Dideoxy sequencing of qPCR amplicons provided species-level identification and phylogenetic validation, including detection of a new species. At three cave sites yielding multiple qPCR detections, 16S metabarcoding enabled species discrimination. By integrating qPCR assay validation, eDNA degradation modelling, hierarchical genetic identification via dideoxy sequencing, and selective NGS metabarcoding, this study presents a strategic framework for applying eDNA methods in subterranean environments.

## INTRODUCTION

The development of environmental DNA (eDNA) techniques has transformed biodiversity monitoring, especially in aquatic environments (Çevik et al., 2025). By detecting trace genetic material shed by organisms into their surroundings, eDNA allows for species identification without direct observation or capture (Ficetola et al., 2008; Thomsen & Willerslev, 2015). DNA fragments from skin, mucous, faeces, or gametes only persist briefly but can be amplified using molecular tools such as quantitative polymerase chain reaction (qPCR) or high-throughput sequencing (Wan et al., 2007; Huang et al., 2010). Over the past two decades, eDNA methods have advanced from single-species detection to ecosystem-level applications (Deiner et al., 2017). Their sensitivity, non-invasive nature, and scalability make them especially effective for monitoring rare or protected taxa that are difficult to capture using traditional methods (Vörös et al., 2017).

Early applications of eDNA focused on detecting individual species in freshwater environments (Jerde et al., 2011; Pilliod et al., 2013; Xia et al., 2021), but advances in eDNA metabarcoding have since enabled the identification of multiple taxa from a single sample (Taberlet et al., 2018; Wang et al., 2021). These advances have expanded the relevance of eDNA across many fields of biological research. However, focus remains predominantly on easily accessible surface environments (Lin et al., 2021). In this context, aquatic subterranean (stygian) habitats such as caves and aquifers remain relatively neglected, despite harbouring biologically interesting species (Culver & Pipan, 2019).

The cyprinid genus *Sinocyclocheilus*, endemic to the karst regions of southwestern China, exemplifies such a lineage (Yang et al., 2016). Comprising over 80 recognised species, with many more likely undiscovered, it inhabits a vast and fragmented karst landscape across Guangxi, Guizhou, and Yunnan provinces (Zhao & Zhang, 2009; Li et al., 2018; Chen et al., 2021; Mao et al., 2022; Liu et al., 2025), covering an area of 218, 025 km² (Liu et al., 2025). Ongoing exploration of these regions continues to uncover new taxa and localities, underscoring both the incompleteness of current inventories and the remarkable biodiversity of China’s subterranean systems (Huang et al., 2023). *Sinocyclocheilus* exhibits stygomorphic traits and varies in eye development, pigmentation, skull ornamentation (horns), fin morphology, neuromast distributions, and cave-specific behaviours (wall-following), representing a rare example of adaptive radiation in subterranean habitats (Mao et al., 2021; Chen et al. 2021; Chen et al. 2022; Mouser et al., 2022). The group has become a key eco-evolutionary model for investigating speciation, adaptation, and biogeographical processes (He et al., 2019; Mao et al., 2025). The “caves as species pumps” proposed for the genus illustrates how repeated cycles of geographical isolation, gene flow, and adaptive divergence produce and sustain diversity (Mao et al., 2022; Mao at al. 2025). Recent genomic research also suggests ongoing introgression and hybridisation among closely related lineages, indicating that speciation within these fishes remains fluid and incomplete (Mao et al., 2025; Liu et al., 2025a, 2025b; 2025c).

Most *Sinocyclocheilus* species are point-endemics, found only in specific caves or springs, making them particularly vulnerable to extinction. Consequently, the entire genus is highly protected (Ma et al., 2023). They face threats from habitat loss, pollution, and changes in water flow caused by agriculture (Zhang et al., 2020). Traditional sampling methods such as netting or electrofishing are often challenging in flooded or narrow passages, and direct observation is limited by poor visibility. In such situations, eDNA provides a powerful, non-invasive tool for detecting and monitoring these fragile populations (Yao et al., 2022; Wang et al., 2024). Collecting water from cave pools or outlets allows for molecular detection without disturbing delicate ecosystems (Heyde et al., 2023). It has been shown that eDNA can be used effectively to detect subterranean species like cave salamanders and groundwater crustaceans (Gorički et al., 2017; Niemiller et al., 2018), but its application to cave fishes remains limited. Thus, *Sinocyclocheilus* serves as an ideal model for developing and testing molecular detection methods (Gao et al., 2024).

Here, we focus on developing and applying a qPCR-dominated assay to detect *Sinocyclocheilus* species, identify new populations of existing and potentially new species, and monitor known populations across southwestern China’s massive karst landscape. We pursued four interconnected objectives: (1) to develop and validate genus-specific qPCR assays targeting the mitochondrial 16S rRNA gene, assessing their specificity and sensitivity against non-target taxa; (2) to examine the effects of temperature and pH on eDNA degradation under controlled, cave-like conditions; (3) to apply the validated assays to environmental water samples from cave and surface habitats, evaluating detection success and spatial variation in eDNA concentrations; and (4) to apply NGS selectively, when qPCR indicates co-occurrence of more than one *Sinocyclocheilus* species at a site.

This strategic integration of molecular approaches links qPCR-based precision (in the context of eDNA in detecting short fragments) with the power of metabarcoding, as needed, to provide an economical and practical framework for documenting, monitoring, and conserving this remarkable yet vulnerable biodiversity across the poorly studied karst ecosystems.

## MATERIALS & METHODS

We aimed to develop, validate, and apply an eDNA assay to detect and monitor fishes of the genus *Sinocyclocheilus*. The research was conducted in three phases (Supplementary Methods S1). The first involved lab-based development of a species-specific quantitative PCR (qPCR) assay targeting the mitochondrial 16S rRNA gene. The second examined eDNA persistence under controlled physicochemical conditions to determine degradation rates across temperature and pH gradients. The third applied the assay to field samples collected from karst habitats in Guangxi, Guizhou, and Yunnan to test its performance under natural conditions. We also employed an eDNA metabarcoding approach to discriminate between species when more than one species was detected by qPCR.

We carried out all molecular procedures involving eDNA before amplification in a dedicated cleanroom facility. Pre-amplification steps, including DNA extraction and standard preparation, were performed in a clean room, while post-amplification steps, including qPCR and sequencing, were performed in another. This physical separation reduced the risk of cross-contamination and preserved the integrity of eDNA analyses.

### Primer and probe design

To develop a reliable, genus-specific eDNA assay for *Sinocyclocheilus*, we designed and tested multiple quantitative PCR (qPCR) primer–probe systems targeting mitochondrial loci (Hutchins et al., 2022). Complete and partial sequences of the 12S and 16S ribosomal RNA (rRNA), cytochrome b (*cytb*), and NADH dehydrogenase subunit 4 (*ND4*) genes were retrieved from the NCBI GenBank database (accessed January 2025). Representative sequences from closely related cyprinid genera (*Cyprinus*, *Carassius*, *Danio*, *Puntius*, and *Gobiocypris*) were also included in the alignments to evaluate potential cross-reactivity.

All sequences were aligned separately by gene region using the MUSCLE algorithm in MEGA v11.0, and conserved *Sinocyclocheilus*-specific regions were identified as candidate targets. Primer and hydrolysis probe sets were designed in PrimerQuest™ (Integrated DNA Technologies, USA) using standard parameters for short amplicons (<150 bp) with melting temperatures between 58 and 62°C and GC content of 40–60% (Supplementary Table 1). We targeted short fragments to facilitate the detection of eDNA that otherwise may be undetectable by metabarcoding. All oligonucleotides were synthesised by Sangon Biotech (Shanghai, China). Each design was assessed for secondary structures and primer–dimer formation using OligoAnalyzer™, and preliminary specificity was examined in silico by BLASTn comparison against the NCBI nucleotide database.

Among the loci tested, primer–probe systems targeting the 12S and 16S rRNA regions exhibited the highest specificity (Beitel et al., 2015). Both probes incorporated a 5′-FAM reporter dye and a 3′-MGB quencher. In contrast, eDNA primers designed from the more variable *cytb* and *ND4* genes, cross-amplified with several non-*Sinocyclocheilus* cyprinids, and were therefore excluded from further validation. Details of locus selection, comparative testing, and *in silico* screening are provided in Supplementary Methods S2.

Initial empirical lab-based evaluations were conducted using genomic DNA extracted from three *Sinocyclocheilus* species representative of the genus (*S. zhenfengensis, S. ronganensis*, and *S. anophthalmus*) and five non-target but closely related cyprinids (as mentioned earlier). Both the 12S and 16S systems successfully amplified only *Sinocyclocheilus* DNA, confirming genus-level specificity (Supplementary Table 1). The 16S rRNA assay, designated JXB-MGB-2, yielded the most consistent amplification and highest efficiency (E = 121%), with no detectable cross-reactivity. Consequently, the JXB-MGB-2 primer-probe system was selected for downstream analyses, including controlled degradation experiments and field validation. The 16S primer region we selected contained a hypervariable region that can distinguish between *Sinocyclocheilus* species.

### DNA extraction and preparation of standards

We obtained fin or muscle tissues from the three *Sinocyclocheilus* species (*S. zhenfengensis*, *S. ronganensis*, and *S. anophthalmus*) and the five non-target cyprinids from ethanol-preserved voucher specimens. Genomic DNA was extracted using the DNeasy Blood & Tissue Kit (Qiagen, Germany) following the manufacturer’s protocol. DNA integrity was examined by agarose gel electrophoresis, and concentrations were measured with a Qubit™ 4.0 fluorometer to ensure accurate quantification for subsequent analyses.

To prepare quantitative standards, the 137-bp *Sinocyclocheilus* 16S rRNA fragment was amplified by conventional PCR and cloned into a pMD-19T vector (Takara, Japan). Recombinant plasmids were transformed into *Escherichia coli* DH5α competent cells and confirmed by dideoxy (Sanger) sequencing. Plasmid DNA was purified using the TIANprep Mini Plasmid Kit (Tiangen, China), quantified fluorometrically, and stored at −20°C until use.

Plasmid copy number was calculated from the measured DNA concentration and molecular weight. Ten-fold serial dilutions ranging from 10L to 10¹ copies per microliter were prepared in sterile nuclease-free water. These dilutions were used to construct the qPCR standard curve and to establish the detection limit of the assay (Detailed transformation and purification steps are provided in Supplementary Methods S1.)

### Quantitative PCR (qPCR) assay

Each quantitative PCR (qPCR) reaction was prepared in a total volume of 20 µL, containing 10 µL of 2× qPCR Master Mix (Yijian Biotech, Suzhou), 0.4 µL of each forward and reverse primer (10 µM), 0.2 µL of probe (10 µM), 2 µL of template DNA, and nuclease-free water to reach the final volume. Amplifications were performed on a LineGene 9600 thermocycler under the following conditions: initial denaturation at 95°C for 30 seconds, followed by 40 cycles of 95°C for 10 seconds, 48°C for 30 seconds, and 72°C for 30 seconds (Mondo et al., 2019). Each run included three no-template controls and three extraction blanks to detect potential contamination.

Serial ten-fold dilutions of the plasmid standards were amplified in triplicate to generate a standard curve relating cycle threshold (Ct) values to logLL-transformed copy numbers. Amplification efficiency (E) was calculated from the slope using the equation E = (10L¹/LLLLL − 1) × 100%. The limit of detection (LOD) was defined as the lowest concentration consistently detected across all replicates, with a standard deviation of less than 0.5 Ct.

Specificity was assessed using DNA extracted from the five non-target closely related cyprinid species (*Cyprinus carpio*, *Danio rerio*, *Puntius semifasciolatus*, *Gobiocypris rarus*, and *Carassius auratus*). Reactions showing no amplification signal (Ct > 40) were considered negative, confirming the absence of cross-reactivity and high target specificity of the primer–probe system (Supplementary Table 2).

### eDNA degradation

A controlled degradation experiment was conducted to examine the persistence of eDNA under different temperature and pH conditions. Nine 100-L polypropylene tanks were prepared, each containing 50 L of deionised water adjusted to pH 4, 7, or 10 using hydrochloric acid or sodium hydroxide (Chen et al., 2023). Temperatures were maintained at 10°C, 18°C, or 26°C, producing nine unique combinations of temperature and pH. Each tank was inoculated with 100 µL of *Sinocyclocheilus* plasmid DNA at a concentration of 10L copies/mL.

At predetermined intervals of 0, 2, 4, 8, 16, 24, 48, 72, 96, 120, 144, 168, and 336 hours following inoculation, 1-L water samples were collected. Three replicate samples and one negative control (DNA-free water) were taken for each condition. Water samples were filtered through 0.45-µm mixed cellulose ester membranes (47 mm diameter), and the filters were immediately stored at −80°C pending DNA extraction.

Following the manufacturer’s protocol, we extract DNA from each filter using the TIANamp DNA Kit (Tiangen, China). Extracts were eluted in 60 µL of TE buffer and quantified by qPCR under the same reaction conditions as previously described. Each sample was analysed in triplicate, and mean values were used for subsequent calculations. Replicates that failed to amplify were recorded as zero copies.

For each treatment, eDNA copy numbers were logLL-transformed and regressed against time to estimate the degradation rate constant (k). Half-lives were derived from these rate constants using the equation t_½_ = ln2/k. The effects of temperature and pH on k were evaluated by two-way analysis of variance (ANOVA) implemented in R version 4.3.0.

### Field sampling and eDNA analysis

We conducted field surveys from March to June 2025 across the massive karst regions in Guangxi, Guizhou, and Yunnan (from 103.79°E to 110.92°E and 24.53°N to 25.21° N). A total of forty-seven sites were sampled, including thirty-seven subterranean cave waters and ten surface streams. These include three caves where multiple species exist (LBY-3, LY-3, SZ-3) (Figure 5). At each site, three 1L water samples were collected in sterile bottles, resulting in nine replicates per location (Sauseng et al., 2025). Environmental parameters, including water temperature, pH, dissolved oxygen, conductivity, total dissolved solids, and salinity, were measured on site using a YSI Professional Plus multiparameter meter (AZ86031, China). To check for possible contamination, 1L blanks of sterile DEPC-treated water were exposed at each location as field controls.

**Figure 1.**
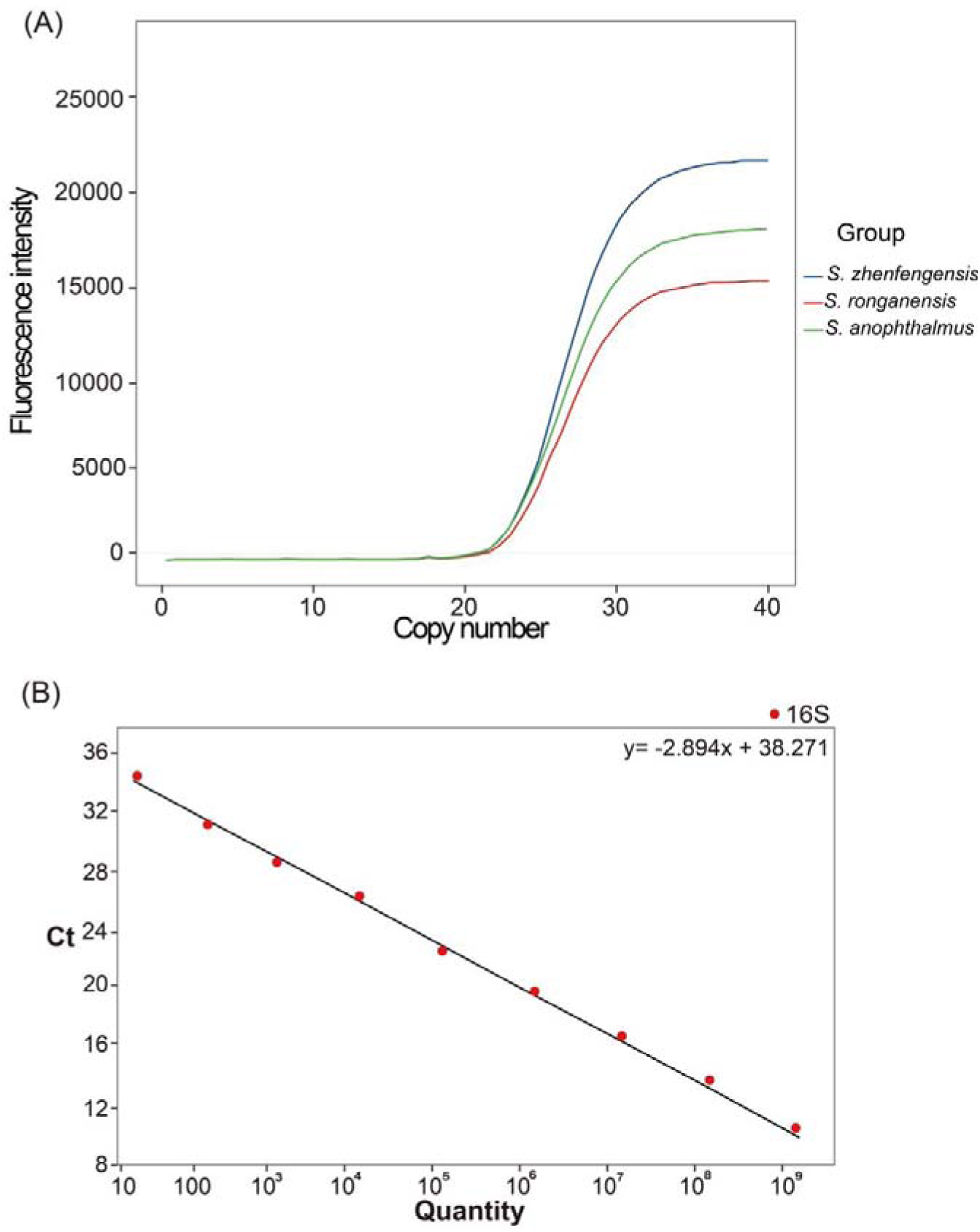
Validation of the *Sinocyclocheilus*-specific qPCR assay. (A) Amplification curves showing successful qPCR detection for three *Sinocyclocheilus* species (*S. zhenfengensis*, *S. ronganensis*, *S. anophthalmus*). Non-target cyprinids and negative controls showed no amplification (Ct > 40). (B) Standard curve generated from ten-fold serial dilutions of plasmid standards, relating Ct value to log10 DNA copy number (R^2^ = 0.995; efficiency = 121.6 %). Both panels confirm high analytical sensitivity and strict genus-level specificity of the JXB-MGB-2 16S rRNA assay.

**Figure 2.**
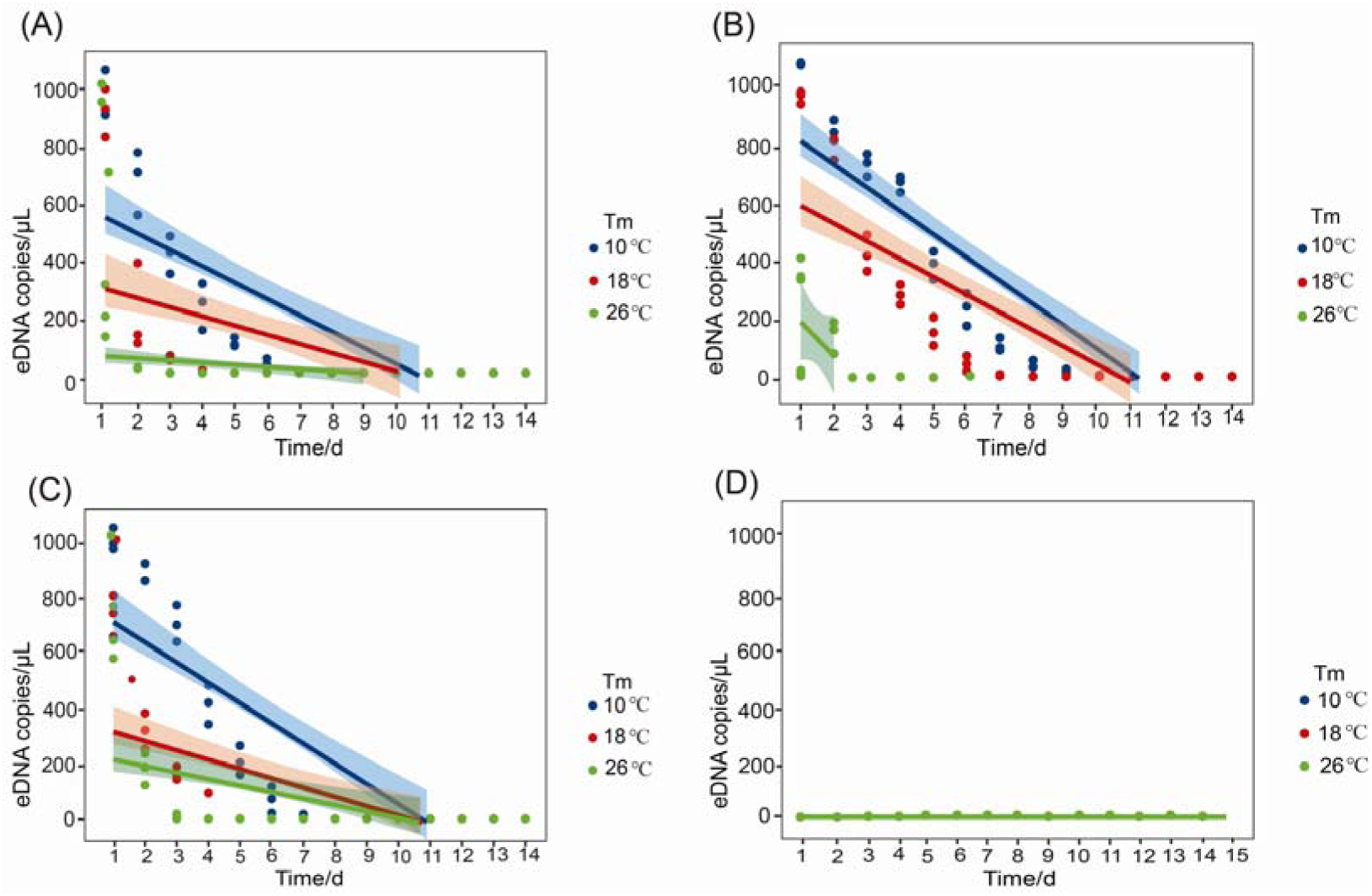
Environmental DNA degradation under varying temperature and pH. Linear regression plots showing changes in *Sinocyclocheilus* eDNA concentration over time under combinations of temperature (10°C, 18°C, 26 °C) and pH (4, 7, 10). Panels (A)–(C) display degradation trajectories for each pH level; panel (D) shows Sterile, enzyme-free water as a negative control (NC), confirming the absence of contamination. Regression models explained more than 90 % of the variance (R² = 0.91–0.97). Two-way ANOVA indicated significant effects of both temperature and pH on degradation rate (p < 0.01).

Samples were transported on ice (<4°C) and filtered within six hours of collection. Filtration was performed under vacuum using 0.45-µm mixed cellulose ester membranes. The filtration apparatus was rinsed with sterile water and sterilised sequentially with 75% ethanol and hypochlorous acid between samples. After filtration, membranes were rolled with sterile forceps, sealed individually in 2-mL microtubes, and stored at −20°C until DNA extraction.

DNA extraction followed the same procedure as the one used in the laboratory assays. Each extraction batch included a negative control. Quantitative PCR assays were conducted in triplicate for every sample, and a sample was considered positive when at least two of three technical replicates exhibited amplification (Ct ≤ 40) with a typical sigmoidal curve. Copy numbers were calculated from the established standard curve and expressed as copies per litre of water.

### Sequencing and bioinformatic analyses

Positive environmental samples were amplified using primer (F: TCGTGCCAGCCACCGCGGTTA, R: TTNTAGAACAGGCTCCTCTAGG). The resulting amplicons were purified using ExoSAP-IT™ and bidirectionally sequenced on an ABI 3730XL platform. Consensus sequences were assembled and aligned in MEGA 11, then compared against *Sinocyclocheilus* reference sequences using BLASTn to confirm taxonomic identity (Including 25 eDNA sequences from *Sinocyclocheilus*, it also encompasses two outgroup species.) (Detailed reference sequence information is provided in the Supplementary Table 4).

Selected samples (where more than one species was detected to be present through qPCR assays) with high eDNA concentrations were subjected to next-generation sequencing on an Illumina MiSeq platform using paired-end 2 × 250 bp reads. Libraries were prepared through a dual-index, two-step PCR protocol (Berry et al. 2011). Raw FASTQ files were processed sequentially to remove adapters with cutadapt, trim low-quality bases with PRINSEQ (minimum Q20), merge paired reads using PEAR, and eliminate chimeras before clustering operational taxonomic units (OTUs) at 97% sequence similarity in USEARCH. Taxonomic assignment was performed against the FishBase reference database, and only sequences showing ≥97% identity were retained for further analysis.

We conducted two phylogenetic analyses based on 16S rRNA. The first analysis involved generating 16S rRNA fragments using the newly designed eDNA primer set for 47 species from reference collections (collected over 7 years of fieldwork), along with 24 putative species from field collections of eDNA-containing water samples. When more than one species was indicated in qPCR results, the sequences were obtained through metabarcoding of the 16S rRNA region. This analysis was primarily designed to assess whether the newly developed qPCR primer performs as expected under field conditions. The second analysis combined a longer fragment of the 16S rRNA gene (approximately 1600 bp) with the eDNA sequences generated in this study to relate field detections to the broader reference database and to infer phylogenetic relationships within a reasonably robust phylogeny.

For each analysis, maximum likelihood trees were inferred in IQ-TREE v1.6.8 with 1000 ultrafast bootstrap replicates, implemented through PhyloSuite. For each 16S dataset, the optimal substitution model was identified with ModelFinder in PhyloSuite using the Bayesian Information Criterion. Bayesian inference was conducted in MrBayes v3.1.2 using the selected substitution models. Two independent runs of 200 million generations were performed, sampling every 20,000 generations and discarding the first 10% as burn-in. Convergence was assessed in Tracer v1.7.1. Single gene trees were also inferred using maximum likelihood. Sequence matches above 97% identity were treated as valid detections (Fig. 3; Fig. 4; Fig. 5A)

**Figure 3.**
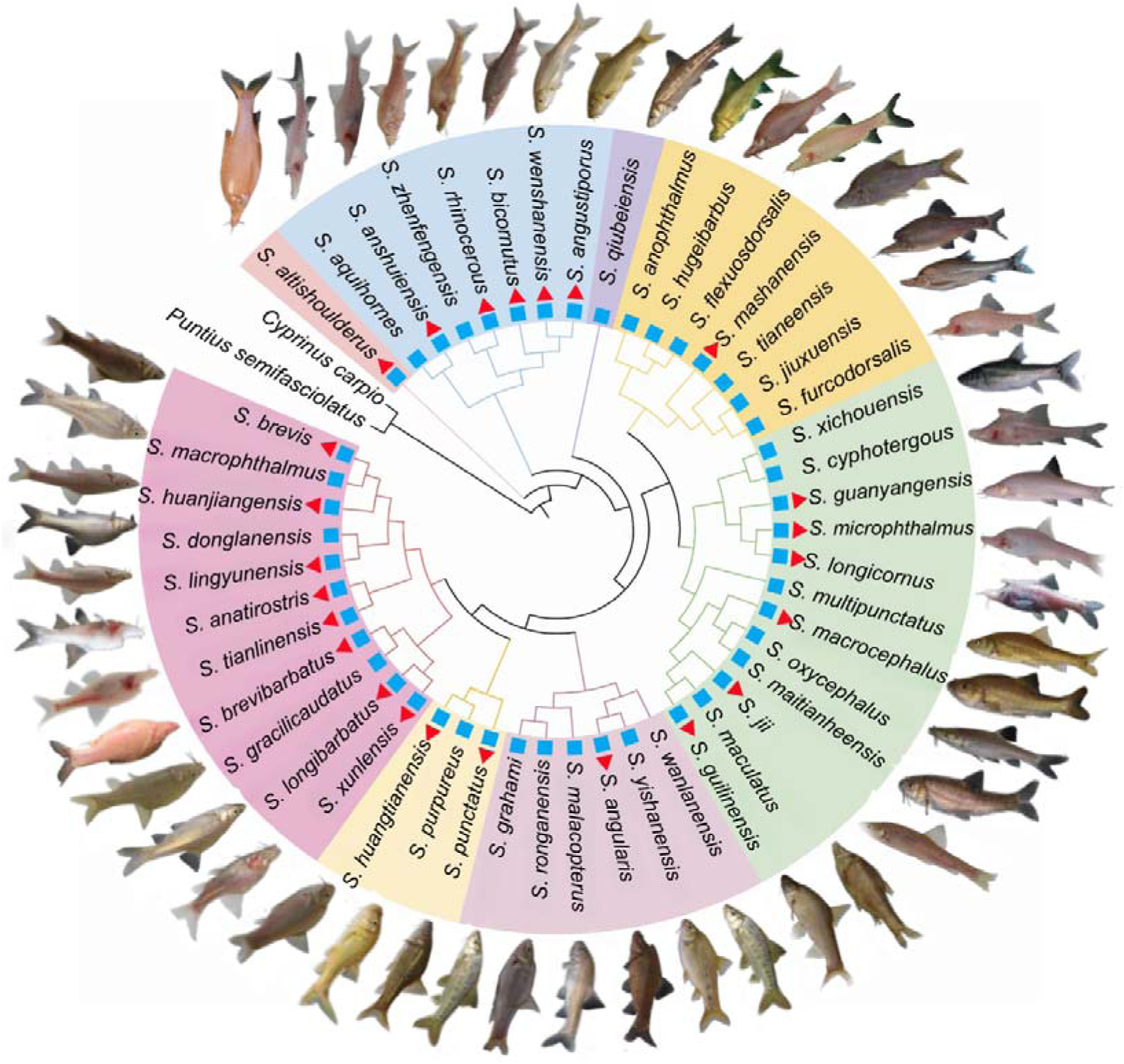
Maximum-likelihood phylogenetic tree constructed from Dideoxy-sequenced 16S qPCR amplicons. Blue squares denote validated samples generated from the eDNA primers, red triangles represent field-detected eDNA. A newly detected species, subsequently described, is also shown in the tree (*S. wanlanensis*). (Detailed construction information is provided in Supplementary Figure 1).

**Figure 4.**
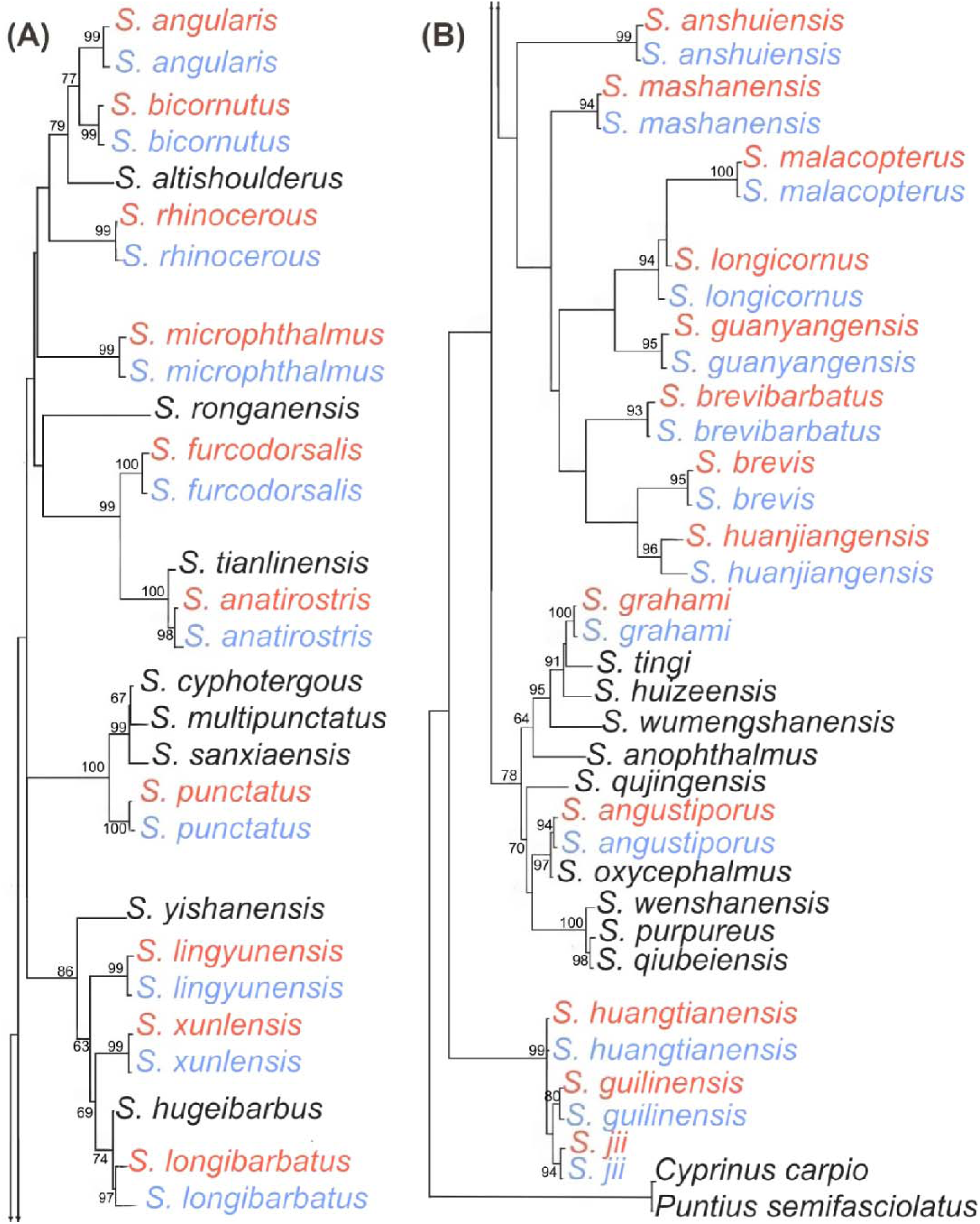
Phylogenetic confirmation of *Sinocyclocheilus* detections from laboratory and field samples. (A & B) Species in red indicate that wild-collected water samples underwent single-generation sequencing and were compared against reference sequences (blue and black) in the database (Detailed reference sequence information is provided in the Supplementary Table 4).

**Figure 5.**
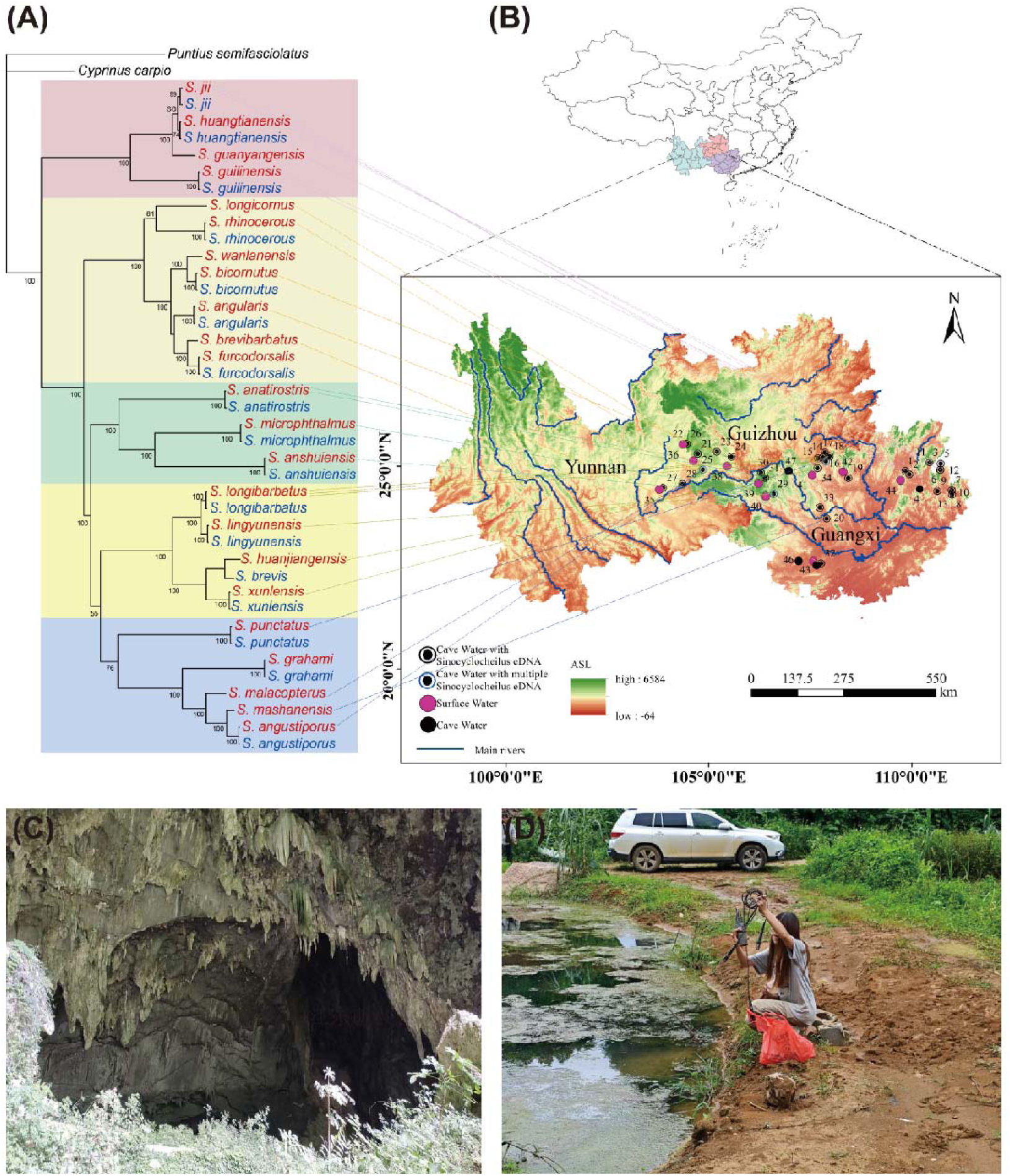
Geographical distribution of sampling sites and eDNA detections across karst regions. Map showing 47 sampling locations (Including 37 cave water sources and 10 surface water sources). (A): Species in red denote eDNA, species in blue denote 16S reference sequences. (Some species lack 16S mitochondrial DNA, such as *S. brevibarbatus*, *S. brevis*, *S. huanjiangensis*, *S. malacopterus*, *S. mashanensis*, *S. longicornus*, *S. guanyangensis*, *S. wanlanensis-*newly described species). (B): Map of sampling point locations across the karst ecosystem. (C): A representative cave water site in Guangxi (106.41E, 24.95N). (D): A representative surface water site close to a cave (106.90E, 22.45N). Geographical coordinates and site details are provided in Supplementary Table 3 (The person in the photo is an author of the paper).

For the first analysis, the resulting tree files were imported into iTOL (Zhou et al., 2023) and visualised using the circular layout, with adjustments to branch, node, and label features for clarity (Fig.3; Fig. 4; Fig. 5A). For the second analysis, where eDNA sequences were placed within a well-supported phylogeny, the ’Branch Metadata Display’ option in iTOL was selected to display bootstraps/metadata.

## RESULTS

### Assay development and validation

The designed primers and probes exclusively amplified *Sinocyclocheilus* DNA. Strong fluorescence signals (Ct 21.5–24.6) were observed from *S. zhenfengensis*, *S. ronganensis*, and *S. anophthalmus*, while no amplification occurred in any non-target cyprinid or negative control (Fig. 1A). Melting-curve analysis confirmed single, species-specific peaks, demonstrating the high specificity of the JXB-MGB-2 probe system for *Sinocyclocheilus*.

Serial dilutions of plasmid standards produced a linear standard curve (Ct = −2.894 logLL[DNA] + 38.271; R² = 0.995) with an amplification efficiency of 121.6% (Fig. 1B). The detection limit was 5 × 10LL ng·µLL¹, equivalent to approximately 20 copies per reaction. Ct values decreased consistently with increasing template concentration, indicating strong quantitative reliability across six orders of magnitude.

### Laboratory eDNA degradation

eDNA concentration declined under all experimental conditions. At neutral pH (7), half-lives were estimated at 68 h (10°C), 45 h (18°C), and 27 h (26°C). Both acidic (pH 4) and alkaline (pH 10) conditions accelerated degradation, reducing half-lives by 35–45%. Regression models explained over 90% of the variance in log-transformed eDNA concentration (R² = 0.91–0.97). Two-way ANOVA confirmed significant effects of both temperature (FL,LL = 45.2, p < .001) and pH (FL,LL = 19.7, p < .01) on degradation rate (Fig. 2).

### Field detection and distribution

Of the 47 environmental samples analysed, 33 yielded positive amplification for *Sinocyclocheilus*, all from cave sites (Fig. 3; Fig. 5). No amplification was detected in any field blank or surface stream samples. Mean Ct values for positive samples ranged from 22.4 to 35.7 (mean ± SD: 28.1 ± 3.7), with estimated eDNA concentrations spanning 3.1 × 10² to 2.6 × 10L copies·LL¹. The highest concentrations occurred in cave pools, while ephemeral streams generally fell below the detection threshold.

Sequences of amplicons approximately 300-360bp in length were selected for sequencing (Fig. 6A). Dideoxy sequencing confirmed that all amplicons matched *Sinocyclocheilus* 16S rRNA sequences with more than 99% identity. BLAST results verified the presence of *S. longibarbatus*, *S. punctatus*, *S. malacopterus*, and *S. rhinocerous* in different localities.

**Figure 6.**
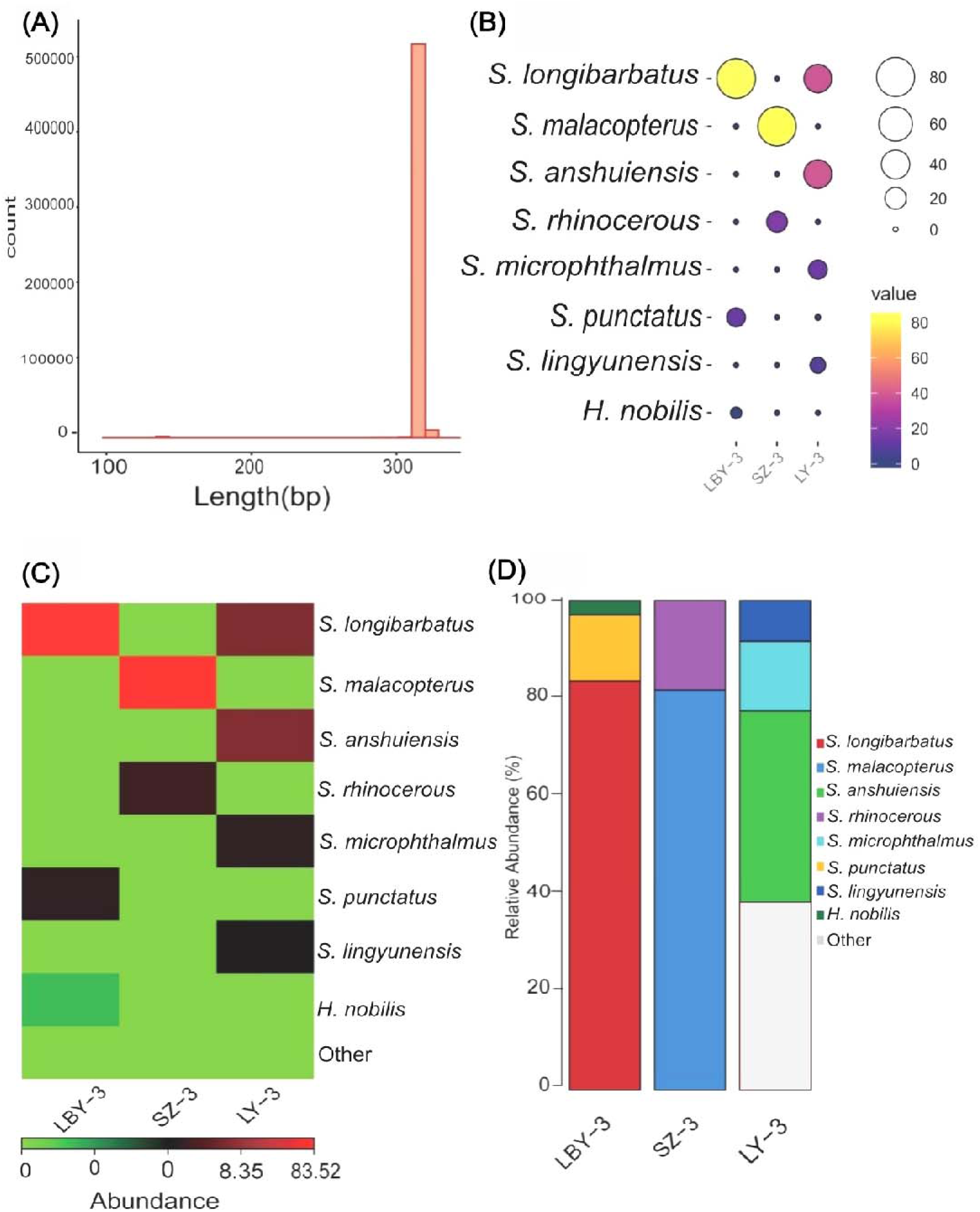
Diversity of *Sinocyclocheilus* assemblages based on next-generation sequencing at the three sites where more than one species was detected using qPCR. (A) Effective sequence length distribution (300 – 360 bp) following quality filtering. (B) Indicates the numerical distribution of seven *Sinocyclocheilus* species (*S. longibarbatus*, *S. malacopterus*, *S. anshuiensis*, *S. rhinocerous*, *S. microphthalmus*, *S. punctatus*, *S. lingyunensis*) across different samples. Darker shades (purple) denote lower values, while lighter shades (yellow) denote higher values. (C) Heatmap of relative abundances for detected species across the sympatric species samples. (D) Bar chart summarising the proportional abundance of species at each site. Collectively, these plots show high spatial heterogeneity and distinct assemblage structures among cave systems.

### 16S metabarcoding of mixed samples

The 16S metabarcoding of the three mixed species cave-water samples (determined to be mixed using qPCR) predicted the following species: LBY-3 contain two species (*S. longibarbatus*, *S. punctatus*); LY-3 has three species (*S. anshuiensis*, *S. microphthalmus*, *S. lingyunensis*); SZ-3 has two species (*S. rhinocerous*, *S. malacopterus*). Heatmap clustering grouped LBY-3 and LY-3, distinct from SZ-3 (Fig. 6B, C). Heatmap clustering grouped LBY-3 and LY-3, distinct from SZ-3 (Fig. 6B, C). Pairwise Bray–Curtis distances supported this structure, showing higher within-group similarity between LBY-3 and LY-3 (Fig. 6D).

### Caves having more than one species

The cave water samples containing more than a single species was subjected t o 16S metabarcoding, which confirmed the sympatric *Sinocyclocheilus* species (Fig.6D; Supplementary Table 5; Supplementary Figure 4; Supplementary Figure 5). Results indicate seven confirmed *Sinocyclocheilus* species were identified a cross the three cave sites. LBY-3 yielded two species of *Sinocyclocheilus* (*S. lo ngibarbatus* and *S. punctatus*), SZ-3 yielded two species (*S. rhinocerous* and *S. malacopterus*), and LY-3 yielded three species (*S. anshuiensis*, *S. microphthalm us*, and *S. lingyunensis*) (Supplementary Figure 6). The distance between the tw o most recently discovered caves exceeds 230 kilometres, and there is no kno wn connection between them (Supplementary Figure 7; Supplementary Figure 8). LY-3 harboured three species, while the other two caves each contained onl y two species. Despite their proximity, no species overlap was observed betwee n the three caves.

## DISCUSSION

Here we developed and validated a quantitative eDNA assay targeting the mitochondrial 16S rRNA gene of the cavefish genus *Sinocyclocheilus*, applying it to controlled experiments and field surveys across the karst ecosystems of southwestern China. Mitochondrial DNA is a single-copy genome but occurs in many copies per cell, making it more readily detectable than nuclear DNA in environmental samples (Uchii et al., 2016). In contrast, primer design for nuclear genes is more complicated, particularly for *Sinocyclocheilus*, which is tetraploid (Mao et al. 2025) and hence presents challenges for locus specificity and consistent amplification.

The assay design proved effective: species identity could be confirmed by sequencing the qPCR amplicons, providing reliable verification in single-species samples. However, this approach is limited when, under some conditions, multiple *Sinocyclocheilus* species co-occur, as mixed templates prevent clear sequence attribution. In such cases, metabarcoding-based techniques are required to resolve community composition and relative diversity, which we did.

Routine cavefish surveys benefit from methods that effectively handle low eDNA levels. Targeted qPCR is well-suited for this, as it more reliably detects low-density material than standard metabarcoding (Blackman et al., 2020; Hii et al., 2024; Yu et al., 2022). It has been shown that qPCR assays maintain sensitivity even when template competition and limited sequencing depth challenge metabarcoding (Johnson et al., 2024). For cavefish, which often exist as isolated populations (low biomass), such sensitivity is crucial for confirming presence, especially in complex environments with dynamic hydrological backgrounds.

Specificity is also high for qPCR assays. qPCR can be thoroughly validated through in silico and empirical tests (Duarte et al., 2023), thereby reducing false positives. Metabarcoding, on the other hand, relies heavily on reference databases and marker performance, both of which remain inconsistent for subterranean fishes (Ushio et al., 2018; Petit-Marty et al., 2023). Many cavefish lineages are still being described, so incomplete reference coverage risks misclassification or no classification at all. Once a qPCR assay is developed for a particular lineage, routine monitoring becomes straightforward, which is ideal for caves where the fish community is usually monospecific and known.

qPCR also provides direct quantification of target copy number, a valuable feature when population sizes are small and demographic trends are difficult to infer from conventional sampling. Measures derived from qPCR often correlate with biomass or abundance (Pont et al., 2022; Baussant, 2022). Metabarcoding can approximate similar quantities when calibrated with internal standards (Ushio et al., 2017; Ledger et al., 2024), but doing so introduces additional steps and remains sensitive to methodological variation. Since many caves require repeated surveys and it is important to detect even rare traces of a single species for conservation, qPCR offers a practical balance of sensitivity, specificity, and precise quantification (Gargan et al., 2021).

The study combined laboratory validation with degradation experiments under controlled conditions and field-based assessments of community composition using quantitative PCR (qPCR) and next-generation sequencing (NGS). Collectively, these analyses demonstrate that the newly developed assay enables reliable and sensitive detection of *Sinocyclocheilus* DNA in both experimental and natural contexts. They further show how environmental parameters influence DNA persistence and reveal spatial variation in community structure among cave systems.

The key outcomes can be summarised into four points. First, the designed primer–probe set achieved strict genus-level specificity, showing no cross-amplification with any of the five tested non-target cyprinids. Second, the assay exhibited high analytical sensitivity, detecting as little as 5 × 10LJLJ ng µLLJ¹ of template DNA, equivalent to roughly 20 copies per reaction. Third, degradation experiments confirmed that eDNA persistence depends heavily on both temperature and pH, with cooler and neutral conditions notably slowing decay. Fourth, application to 41 field sites yielded 28 positive detections, all within cave habitats, and revealed significant geographical heterogeneity in *Sinocyclocheilus* assemblages.

Together, these findings confirm that eDNA methods can be effectively adapted to monitor cryptic cave fishes (Liu Y., et al., 2025), extending the demonstrated utility of the technique from surface aquatic systems to subterranean karst environments. They also emphasise that, with proper validation, eDNA can support both presence–absence surveys and community-level inference for taxa previously considered inaccessible to non-invasive sampling.

### Assay performance and specificity

The success of any eDNA study heavily relies on assay specificity and sensitivity (Goldberg et al., 2016; Furlan et al., 2020). The designed primer–probe system demonstrated complete exclusivity for *Sinocyclocheilus*, with no amplification of related cyprinids. This is a vital development because the genus coexists with morphologically similar taxa in many karst streams, where conventional visual identification can be challenging. The lack of cross-amplification agrees with other studies using short mitochondrial fragments for genus-level assays, such as those developed for *Phoxinus* (Evans et al., 2017) and *Oncorhynchus* (Laramie et al., 2015).

The observed amplification efficiency (121%) remains within acceptable limits for quantitative estimation (Bustin et al., 2009). Such increased efficiency probably results from minor deviations in baseline fluorescence or amplicon size; similar patterns have been observed in high-copy mitochondrial assays (Wilcox et al., 2013). The linearity over six orders of magnitude (R² = 0.995) indicates that the assay offers reliable quantitative performance, supporting its use for semi-quantitative comparisons of eDNA abundance across samples.

An important methodological enhancement is the use of a hydrolysis (TaqMan) probe instead of SYBR Green detection. Although probe-based assays are more expensive, they significantly reduce the risk of non-specific signals and are therefore well-suited for field surveys in complex microbial backgrounds (Jane et al., 2015). Given the high potential for contamination in cave systems, this design choice probably contributed to the absence of false positives in negative controls.

During assay development, we also examined multiple mitochondrial loci to identify the most effective target for *Sinocyclocheilus* eDNA detection. The final 16S assay selected for validation (JXB-MGB-2) includes a hypervariable segment (when considering between genera) that, however, enables differentiation among species within the *Sinocyclocheilus* genus, thereby allowing both detection and initial identification. This makes it especially useful for locating undocumented or isolated populations where morphological sampling is not possible. The additional assays eveloped from 12S and 16S rRNA regions also demonstrated strong genus specificity and similar sensitivity, and they will be used in future comparative tests to improve resolution and extend applicability across the group. Conversely, primer-probe sets targeting *cytB* and *ND4*, despite their higher nucleotide variability, consistently amplified non-target cyprinids, confirming that these protein-coding loci exhibit limited selectivity for this genus in eDNA contexts. Overall, these results underscore the suitability of rRNA loci for sensitive and dependable detection, offering a practical basis for future multiplex or multi-locus applications aimed at resolving cryptic diversity within *Sinocyclocheilus* (see Supplementary Methods S1).

### eDNA degradation and environmental factors

The degradation experiment showed a consistent relationship between eDNA persistence, temperature, and pH. At 10°C and pH 7, DNA remained detectable for up to 14 days, whereas at 26°C and at pH 4 or 10, signals declined to near zero within 4 days. These patterns are in agreement with previous studies indicating that higher temperatures and deviations from neutrality increase hydrolysis and microbial activity, thereby accelerating decay (Barnes and Turner, 2016; Strickler et al., 2015). The rates observed here are comparable to those reported for salmonids (Balasingham et al., 2018), and the effects of temperature and pH resemble findings for amphibians and carp (Takahara et al., 2012; Strickler et al., 2015).

The estimated half-lives (27–68 hours) closely match those seen in similar aquatic environments: Thomsen et al. (2012) reported amphibian DNA persistence for 2–6 days at 15–20°C, while Balasingham et al. (2017) found residual salmon DNA detectable for up to 11 hours in a river system. Tatsuya Saito et al. (2021) reported that fish DNA can persist in pond water for 3–5 days. Jo T S. A et al. (2023) documented that water temperature regulates eDNA release by influencing fish metabolism: while the optimal growth temperature for Japanese jack mackerel is approximately 20°C, at 28°C—though above the optimum—increased metabolic rates (such as respiration and excretion) and physiological stress levels still enhance eDNA release from sources including skin cells and mucus (Luo et al., 2023). Qian et al. (2022) documented that eDNA degradation rates accelerate significantly with rising temperatures, exhibiting an exponential decline trend. The longer persistence observed under our neutral, low-temperature conditions likely reflects lower microbial loads and reduced UV exposure in controlled tanks.

These degradation dynamics have important practical implications for interpreting field detections. In cave systems with stable light and temperature and limited microbial activity, eDNA may persist longer than in surface streams. This increased persistence improves detectability but may also reduce the temporal resolution of presence data: a positive detection might indicate recent occupation rather than immediate presence. Future research can directly measure degradation in representative caves to improve the interpretation of eDNA signals.

### Field detections and distribution patterns

The field survey revealed a clear distinction between cave and surface habitats. All positive detections occurred in caves, while none were obtained from surface streams. This result supports ecological expectations: most *Sinocyclocheilus* species are obligate cave dwellers, with only a few surface or epigean forms (Zhao and Zhang, 2020). The absence of signals in surface waters reinforces the assay’s specificity and indicates minimal environmental contamination or DNA transport from cave sources.

The variation in eDNA concentration among cave sites likely reflects differences in population density, hydrological flow, and microhabitat stability. Sites such as LBY-3, which showed the highest eDNA concentrations, are characterised by stagnant pools with little inflow or outflow. Such environments promote DNA accumulation and longer persistence. Conversely, lower concentrations at SZ-3 probably result from active flow, which enhances dilution and downstream transport. Similar spatial gradients in eDNA concentration related to hydrological flow have been documented for fish (Deiner and Altermatt, 2014) and amphibians (Pilliod et al., 2014).

Interestingly, the absence of detectable DNA in some known cave systems may not necessarily mean the absence of fish. Given the low biomass typical of cave populations, stochastic sampling effects are likely. Each 1 L water sample represents a tiny fraction of the total cave water volume; even with triplicate sampling, the detection probability may remain below 1 (Schmelzle and Kinziger, 2016). Using occupancy modelling frameworks (MacKenzie et al., 2002) in future analyses would allow explicit estimation of detection probability and help differentiate between true absence and non-detection.

### Methodological considerations and potential biases

Rigorous contamination control is essential for the reliability of eDNA detection (Goldberg et al., 2016). The absence of amplification in all field blanks and negative controls confirms that contamination was minimised. However, the possibility of trace carryover between samples cannot be ruled out. Field blanks were filtered alongside environmental samples, but air- or aerosol-borne DNA could still introduce low-level contamination.

Though qPCR provides relative measures of DNA concentration, converting these into biomass or abundance remains difficult. Variability in shedding rates, hydrodynamics, and DNA degradation complicates direct links between copy number and population size (Takahara et al., 2012). The high variation in Ct values among replicate samples at the same site illustrates this challenge. Instead of inferring absolute abundance, eDNA concentrations should be viewed as indicators of relative activity or habitat use intensity.

While the 16S rRNA marker allowed genus-level detection and, in many cases, species-level discrimination, some *Sinocyclocheilus* species share highly similar mitochondrial sequences. Using additional loci (e.g., COI, cyt b, or nuclear markers) could improve taxonomic resolution (Gorički et al., 2017). Alternatively, genomic methods such as eDNA capture or shotgun metagenomics may permit finer-scale identification, though at higher cost (Sigsgaard et al., 2020).

The NGS data produced abundant reads, but a small proportion of low-frequency OTUs might be sequencing artefacts or result from tag-jumping. The 97% clustering threshold balances sensitivity with error control but may underestimate intraspecific diversity. Future analyses could employ denoising algorithms (e.g., DADA2 or UNOISE3) to recover exact sequence variants and better distinguish closely related haplotypes (Callahan et al., 2016).

### Conservation implications

Cave ecosystems pose significant challenges for biological surveys due to limited access, low light levels, and the often protected status of fauna (Guzik et al., 2024). Traditional sampling methods, such as netting or electrofishing, are often impractical or unethical in these complex subterranean environments. The non-invasive nature of eDNA analysis offers a transformative tool for monitoring such systems. This study demonstrates that short fragments of mitochondrial DNA can be recovered and identified even from small volumes of cave water, enabling species inventories without direct sampling of organisms.

The ability to detect rare species has clear conservation implications. Many *Sinocyclocheilus* species are classified as critically endangered because of habitat loss, groundwater extraction, and pollution (Zhang et al., 2020). Regular eDNA surveys could provide early warning of local extirpations or colonisations, supporting adaptive management of karst aquifers.

The spatial distribution of eDNA detections may also indicate hydrological connectivity between cave systems. Since DNA molecules can be transported with groundwater flow, detecting shared haplotypes in separate caves might signify physical linkage. When combined with hydrological tracing, such inferences could significantly improve our understanding of subterranean drainage networks (Deiner et al., 2017).

eDNA offers high sensitivity, and it can complement traditional surveys. Physical specimens remain crucial for morphological verification and voucher deposition when new species are describe. Nevertheless, eDNA can guide the discovery of new species and monitoring efforts when specimens are not needed, reducing disturbance and enhancing efficiency. Over time, integrating such datasets could support population models that combine genetic, environmental, and hydrological data to predict species distributions under changing climatic conditions.

### Comparison with previous eDNA studies

The patterns of eDNA dynamics agree with previous studies on fish in both controlled and natural systems. For example, Sigsgaard et al. (2016) achieved similar sensitivity when detecting whale sharks from seawater samples, with eDNA sequences matching tissue haplotypes. Likewise, Dugal et al. (2022) showed that individual whale sharks could be “haplotyped” from eDNA, attaining 100% match rates between tissue and environmental samples.

While this study shows that eDNA performs well in cave settings, several constraints require attention in future work. Our sampling took place at only one point in time, although seasonal changes in hydrology and biological activity are likely to affect DNA persistence. Repeated sampling across wet and dry periods would help to better characterise these temporal patterns (White et al., 2018). We also did not estimate detection probability, but the use of occupancy models in future studies would enable the assessment of false negatives and provide more solid statistical support (MacKenzie et al., 2002).

Relying solely on mitochondrial markers limits interpretation to maternal lineages. Nuclear loci could offer deeper insights into population structure (Littlefair et al., 2022), but their use in Sinocyclocheilus remains challenging due to its tetraploid genome (Mao et al. 2025). Laboratory findings concerning DNA degradation should also be approached with caution, as conditions in subterranean environments can accelerate DNA breakdown. Microbial biofilms and mineral adsorption, for instance, can enhance the loss of genetic material (Zhou et al., 2022).

The integration of qPCR and NGS also demonstrates how eDNA studies are moving from monitoring individual species to assessing entire ecosystems (Niemiller et al., 2018). Quantitative assays provide reliable presence-absence and relative abundance data, while metabarcoding reveals community structure and potential species interactions (Alter et al., 2022). In this way, the approach mirrors the shift seen in marine systems from merely detecting species to exploring population genetics, as illustrated by Dugal et al. (2022) and Sigsgaard et al. (2020). Applying this to *Sinocyclocheilus* therefore signifies a conceptual breakthrough: using eDNA not just for monitoring but also for investigating patterns of endemism and evolutionary divergence in subterranean lineages (Gorički et al., 2018).

## CONCLUSIONS

The development and validation of a genus-specific qPCR assay for *Sinocyclocheilus*, along with controlled degradation experiments and extensive field sampling, demonstrate that eDNA can be a reliable tool for surveying cavefish diversity. The method provides a non-invasive, repeatable, and sensitive alternative to traditional sampling in fragile karst systems.

Environmental conditions significantly influence DNA persistence, with cooler and neutral waters supporting longer detection periods. Field data show notable spatial variation in *Sinocyclocheilus* populations, emphasising the ecological uniqueness of each cave system. These findings not only confirm eDNA as an effective monitoring tool for subterranean fauna but also underline its potential to guide conservation strategies for one of China’s most distinctive and endangered fish radiations.

Future research should increase temporal and spatial replication, incorporate multi-marker or genomic approaches, and connect molecular signals with hydrological and ecological models. Through such efforts, eDNA analysis can become a fundamental method in the study and preservation of subterranean biodiversity.

## Acknowledgements

We are grateful to the following individuals: Zhou Shipeng, Chen Bing, and Fu Chenghai and Zhou Jiajun for their assistance with field sampling; and the Departments of Agriculture and Rural Affairs of Guangxi, Yunnan, and Guizhou for their support during fieldwork.

## Author Contributions

MM, YMP, TRM, YWL, MRP, and JY conceived the study and designed the methodology. YMP, SJY, RJC, YWL, TRM, MM, and JY conducted fieldwork and collated data. YMP, YWL, MM, and SJY performed the formal analyses. MM, YMP and YWL first drafted the manuscript. MM, MRP, DS and JY supervised. MM and JY secured research funding. YMP, MM, SJY, TRM, and YWL produced figures and tables. All authors reviewed and revised the draft. All authors read and approved the final manuscript.

## Funding sources

(1) National Natural Science Foundation of China (Project #3226033) for *Sinocyclocheilus* eDNA research; (2) Guangxi Science and Technology Base and Talent Special Project (AD25069066); (3) MRP was funded by a scholarship from National Council for Scientific and Technological Development - CNPq (grant no. 303491/2024-8). These funding bodies played no role whatsoever in the research design, data collection, analysis and interpretation, or manuscript preparation.

## Availability of data and materials

All data generated or analysed during this study or the sources of data are included in this published article. The working primer sequences and DNA data generated will be provided upon acceptance for formal publication.

## Ethics approval and consent to participate

The treatment of experimental animals in this study fully complies with the Chinese Animal Welfare Law (GB/T 35892-2018). Specimens were collected with permission from the Guangxi, Guizhou and Yunnan Provincial governments. All animal protocols in this study were reviewed and approved by the Research Animals Ethics Committee of Guangxi University (Approval No. GXU-2024-278).

## Competing interests

The authors declare that they have no competing interests.

## Notes

### Competing Interest Statement

The authors have declared no competing interest.

